# Genetically engineered resistance to bufotoxin in marsupial *ATP1A1*

**DOI:** 10.1101/2024.05.07.591791

**Authors:** Pierre Ibri, Gerard Tarulli, Sara Ord, Andrew J. Pask, Stephen R. Frankenberg

## Abstract

The introduction of the bufotoxin-secreting cane toad (*Rhinella marina*) to Queensland in 1935 has had a devastating impact on wildlife in the Australian tropics. Having evolved for millions of years in the absence of cane toads or other bufotoxin-secreting organisms, many of the Australia’s native predators that include cane toads in their diet suffered large population declines following cane toad invasion to their habitat. One marsupial species, the northern quoll (*Dasyurus hallucatus*), is now classified as endangered (IUCN Red List) largely due to bufotoxin ingestion. This study aimed to introduce bufotoxin resistance into a marsupial cell line by editing part of the *ATP1A1* gene encoding the extracellular H1-H2 domain – the binding target of bufotoxin. To this end, CRISPR prime editing was used to replace the part of the wildtype *ATP1A1* gene encoding the H1-H2 domain in fibroblasts of a related marsupial model, the fat-tailed dunnart (*Sminthopsis crassicaudata*), with modifications known to be associated with bufotoxin resistance. The genetically modified cell population showed a >45-fold increase in resistance to bufalin (an active component of bufotoxin) compared to wild type. This study provides a proof of concept towards engineering genetic resistance in the northern quoll to halt or even reverse its current population decline.

## Introduction

Invasive species pose an ongoing threat to the world’s biodiversity (Middleton, 2008). The introduction of a species to a new habitat can negatively impact native flora and fauna populations (Middleton, 2008), which can be even more severe on islands (Fritts & Rodda, 1998). Island-living species lack interaction with foreign species during their evolution, making them particularly prone to introduced threats from other parts of the world (Fritts & Rodda, 1998). In Australia, an island continent, many invasive species have been introduced, either accidently or deliberately, since European settlement in 1788 (Department of Climate Change, Energy, Environment, and Water [DCCEEW], 2022). Home to hundreds of thousands of species with high levels of endemism, Australia is one of the most biodiverse countries on the globe (Woinarski et al., 2015). However, introduction of invasive species in addition to habitat loss and altered bushfire regimes have driven many of its species to extinction and put many others at the risk of going extinct (Woinarski et al., 2015). Since European colonisation of Australia, around one hundred confirmed species have gone extinct, including 34 mammals (more than any other country), with more than 1700 others currently classified as threatened (DCCEEW, 2021; Woinarski et al., 2019; Woinarski et al., 2015)

The cane toad (*Rhinella marina*) was introduced to Australia from Hawaii in 1935 to control beetles in sugar cane fields in Queensland (Lever, 2001). They are large toxic anurans, with a native range that stretches from the northern parts of Latin America to southern Texas in the United States (Lever, 2001). Like other members of the family Bufonidae, cane toads secret bufotoxins from their parotid glands as a mechanism of defence when being attacked by a predator (Lever, 2001). The cane toad’s highly potent toxin contains a cardiac glycoside called bufadienolide that targets the sodium/potassium ATPase enzyme present on the cellular membrane of animal cells (Lever, 2001; Plumlee, 2004; Rodríguez et al., 2017). The inhibition of the sodium/potassium ion transfer across the cellular membrane alters its resting potential, leading to impairment of electrical conduction in excitable cells such as cardiac muscles (Plumlee, 2004). Since their introduction to previously toad-free continent, cane toads have managed almost all areas of tropical Australia (Kearney et al., 2008). With no previous exposure to any bufotoxin-secreting species, many Australian predators have seen drastic declines in their numbers due to attempted predation on cane toads, such as freshwater crocodiles, varanid lizards, and northern quolls (Doody et al., 2006; Jolly et al., 2015; Letnic et al., 2008; Moore et al., 2019; Phillips et al., 2010; Shine, 2010; Ujvari & Madsen, 2009; Ujvari et al., 2013b). Nonetheless, some snake, rat, and frog species have shown rapid adaptation to cane toads (Greenlees et al., 2010; Parrott et al., 2020; Phillips & Shine, 2004, 2006).

A previous study by (Ujvari et al., 2013a) showed that small variations in the amino acid sequence of the extracellular H1-H2 domain of ATP1A3 (the alpha 3 isomer of the Na^+^/K^+^ pump) are responsible for the high bufotoxin tolerance among Asian/African varanids (bufotoxin-resistant) compared to Australian varanids (bufotoxin-susceptible). Further DNA analysis revealed similar patterns causing resistance to bufotoxins in other species that co-evolved with bufotoxin-secreting organisms (Ujvari et al., 2015). These patterns seem to be consistent among species of the same group of animals and vary slightly between the different animal groups (Ujvari et al., 2015). Bufotoxin-resistant mammals, for example, have these modifications in the H1-H2 domain-encoding sequence of the *ATP1A1* gene compared to *ATP1A3* as seen among reptiles (Ujvari et al., 2015).

There has been limited success in attempts to control the cane toad invasion, which is predicted to reach more regions of Australia (Kearney et al., 2008; Shanmuganathan et al., 2010; Shine et al., 2018). Therefore, more efforts have been directed toward minimizing their impact on Australia’s ecosystem (Indigo et al., 2018; Kelly & Phillips, 2019; O’Donnell et al., 2010). Here, we report the genetic engineering of bufotoxin resistance in the *ATP1A1* gene of fibroblast cells of the fat-tailed dunnart, a small, carnivorous marsupial in the same family (Dasyuridae) family as the northern quolls). Our result establishes a proof-of-principle for potentially engineering bufotoxin resistance in the northern quoll to reverse its current population decline.

## Methods and Materials

### Dunnart Cell Line Derivation and Culture

The fibroblast primary cell line used in the study was derived from a one-day-old fat-tailed dunnart pouch young. Fibroblast cells were cultured in a medium containing Dulbecco’s Modified Eagle Medium (DMEM) (ThermoFisher Scientific), 10% Fetal Bovine Serum (FBS) (ThermoFisher Scientific), and 1× Antibiotic-Antimycotic (ThermoFisher Scientific). Medium was changed every two days and cells were passaged as required.

### Introducing the Genetic Edits

Prime editing guide RNAs (pegRNAs) used in this study were designed using pegFinder online tool (Chow et al., 2021) and then ordered as gBlocks from Integrated DNA Technologies (IDT). In-Fusion cloning (TakaraBio) was used to separately clone the different pegRNAs into the piggyBac plasmid vector (Addgene #173222), which contains the prime editor (PE2), and a puro/ΔTK selection system (Wolff et al., 2021) as per the manufacturer’s instructions. Lipofectamine 3000 (ThermoFisher Scientific) was used to chemically transfect dunnart fibroblasts with 2.5 μg of plasmid DNA mix (including the pegRNA-cloned PE2 plasmid and piggyBac transposase plasmid at a ratio 9:1) per one million cells. On day 3 post-transfection, medium was changed, and the cells were subjected to puromycin selection (4 μg/mL) for two weeks, followed by bufalin selection (0.1 μM) for a further 2 weeks to obtain homogeneous cell populations. The successful introduction of the different genetic edits was confirmed by Sanger sequencing at the Australian Genome Research Facility (AGRF).

### Cytotoxicity Assay

Wild type and edited cell populations were seeded in 96-well plates (10,000 cells/well) and allowed to proliferate in the absence of bufalin until reaching 50% confluency. The medium was then changed and the cells subjected to different concentrations of bufalin for 24 hours. Finally, the water-soluble tetrazolium 1 (WST-1) (Cayman Chemicals) assay was used as per the manufacturer’s instructions, and the absorbance was read at 440 nm wavelength two hours later using a microplate reader (BMG LabTech). All graphs and statistical results were produced using Graph Pad Prism.

## Results

The DNA sequence data of the genetically modified cell population (Fig. 1) confirmed the successful introduction of the targeted edits into one of the two copies of the *ATP1A1* gene (equal signal intensity of the wildtype and introduced bases at the targeted sites).

**Figure 1:**
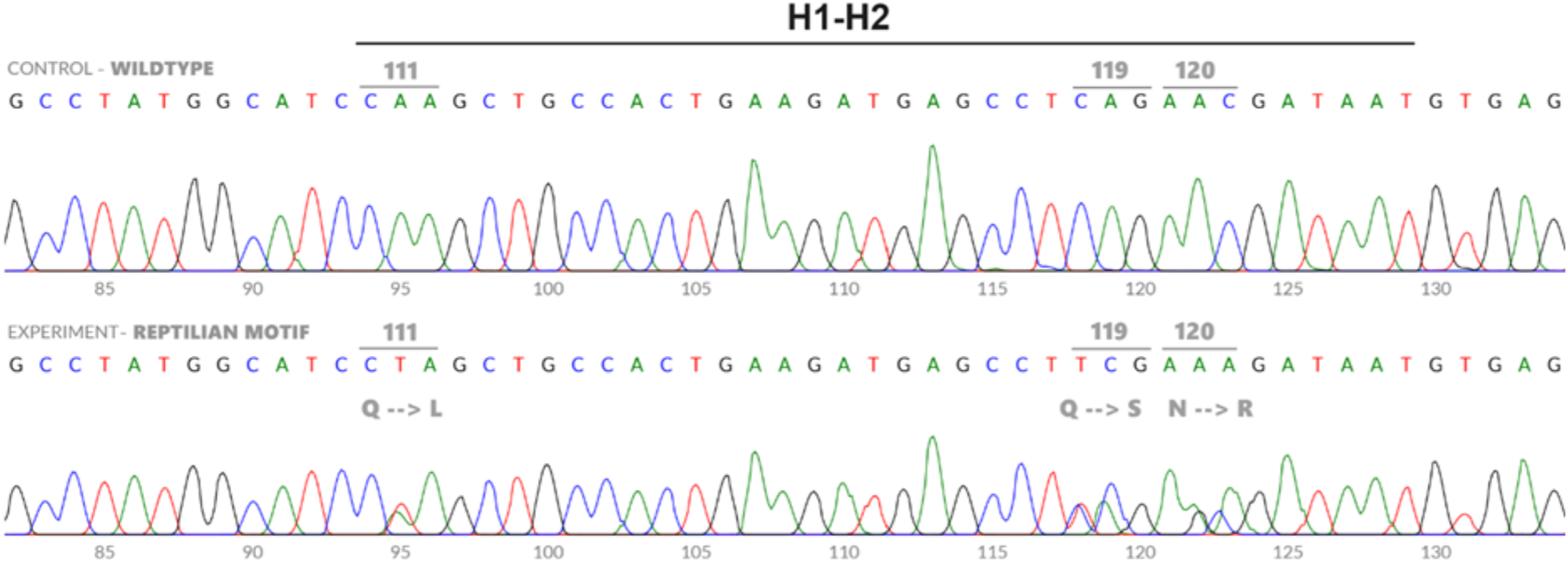
DNA sequences of the H1-H2 domain of the *ATP1A1* gene of the wildtype dunnart cell line (top), and the genetically modified dunnart cell line with the varanid bufotoxin resistance edits introduced into positions 111, 119, and 120 (bottom). DNA sequences were viewed using the online tool DECODR (Bloh et al., 2021).

The wildtype and the CRISPR-edited cell populations showed similar cell proliferation rates at the lowest bufalin concentrations 0.001 uM, 0.003 uM, 0.006 uM (*t*_*160*_ *= 1*.*38, p-value = 0*.*17, t*_*160*_ *= 0*.*70, p-value = 0*.*48, t*_*160*_ *= 0*.*41, p-value = 0*.*69* respectively) (Fig. 2) and were significantly different at all other concentrations.

**Figure 2:**
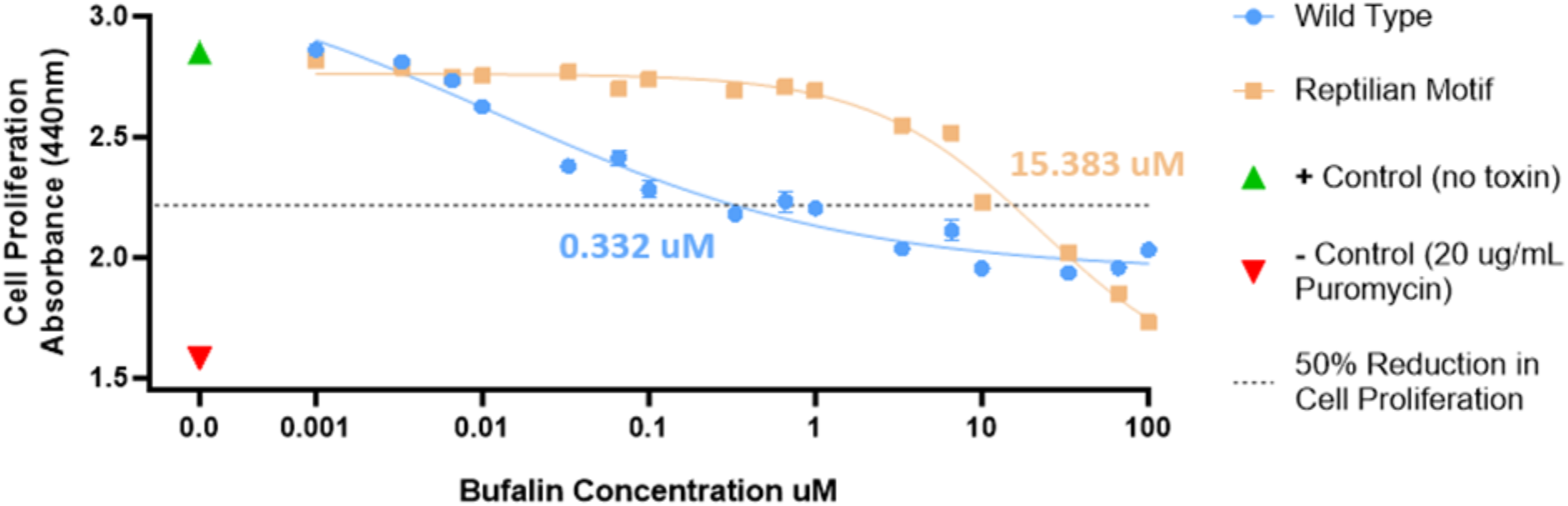
Cell proliferation rate (measured as absorbance at 440 nm wavelength using WST-1 assay) of the wildtype (non-transfected fat-tailed dunnart fibroblasts) and the CRISPR-edited fat-tailed dunnart fibroblasts (with the varanid bufotoxin resistance edits introduced into the H1-H2 domain of the *ATP1A1* gene) against different bufalin concentrations, with positive control (non-transfected fat-tailed dunnart fibroblasts with no exposure to bufalin) and negative control (non-transfected fat-tailed dunnart fibroblasts exposed to 20 µg\mL puromycin). Error bars represent standard error of mean (SEM). The dotted line represents 50% reduction in cell proliferation rate between the positive and negative controls. Non-linear regression curves were fitted to the mean cell proliferation values at each bufalin treatment of the wildtype (*R*^*2*^ *= 0*.*93*) and CRISPR-edited cell populations (*R*^*2*^ *= 0*.*97*).

50% reduction in cell proliferation occurred in wildtype cells at a bufalin concentration of 0.332 µM, and in the edited cells at 15.383 µM, showing over 45-fold increase in bufalin tolerance compared to non-transfected cells.

## Discussion

The Na^+^/K^+^ ATPase enzyme plays an important role in many cellular functions, such as maintaining the resting potential of the cellular membrane and regulating the levels of sodium and potassium ions in the cytoplasm (Clausen et al., 2017). Many species produce toxins that target Na^+^/K^+^ pump for defence against predators (Clausen et al., 2017). In addition to members of the family Bufonidae, to which the cane toad belongs, multiple plant families secret similar cardiac steroids that inhibit the Na^+^/K^+^ pump as a mechanism of defence (Clausen et al., 2017). Resistance to these cardiac steroids is also common among animals that co-evolved with these bufotoxin-secreting organisms and adapted to feeding on them (Ujvari et al, 2015). The differences observed between species in the acquired genetic changes that cause bufotoxin resistance, such as the amino acids on either side of the H1-H2 domain and the specific isomer of the alpha subunit containing these amino acids (Ujvari et al, 2015), suggests that some patterns are more favourable than others. Here, we chose to target the *ATP1A1* gene since it tends to have acquired resistance-associated modifications among the bufotoxin-resistant mammals included in the comparison (Ujvari et al, 2015)

The introduction of the varanid bufotoxin-resistance motif (Leucine in 111, Serine in 119, and Arginine in 120 positions of the H1-H2 domain) into *ATP1A1* of fat-tailed dunnart fibroblasts, led to >45-fold increase in bufalin tolerance compared to wild type (untransfected dunnart fibroblasts). This increase in resistance to bufalin, albeit significant, is far less than the 3000 times increase in toxin resistance among the transfected HEK293 cells in a previous study (Ujvari et al, 2013a). This large difference in results is unlikely to be due to the different cell lines used – fat-tailed dunnart fibroblasts in this study versus human embryonic kidney HEK293 (Ujvari et al, 2013a). Rather, it is likely due to the different strategies adopted in each study. Here, we used CRISPR Prime Editing to precisely introduce the edits into the endogenous *ATP1A1* gene and the result was a heterozygous cell population with one resistant allele and one wild-type allele (Fig. 1). On the other hand, Ujvari et al (2013a) transfected the HEK293 cells with a plasmid that contains a modified version of the human *ATP1A1* gene that contains the targeted bases of the H1-H2 domain, preceded by a strong promoter (CMV). The later method results in multiple copies of the exogenous modified *ATP1A1* transgene being introduced to the target cells, likely resulting in a very high ratio of resistant to wildtype ATP1A1 protein. This difference in the results between the two studies suggests that the level of bufotoxin resistance is dependent on the level of the bufotoxin-resistant version of the ATP1A1 protein. Therefore, a homozygous cell population is likely to show higher bufotoxin resistance.

## Acknowledgements

The authors thank the support of Colossal Biosciences for assistance in CRISPR priming editing methods and Territory Wildlife Park, Dr Richard Mollard, Dr Emily Scicluna and Dr Mitzi Klein for assistance in tissue collections. This study was supported by the Hermon Slade Foundation, the Australian Research Council, the Wilson Family Trust, and Colossal Biosciences.

